# SCNT: An R Package for Data Analysis and Visualization of Single-Cell and Spatial Transcriptomics

**DOI:** 10.1101/2025.06.02.657520

**Authors:** Jianbo Qing, Jialu Wu, Yafeng Li, Junnan Wu

## Abstract

**Background:** The emergence of single-cell (SC) and spatial transcriptomics (ST) has revolutionized our understanding of gene expression dynamics in complex tissues. However, it also presents challenges for data analysis and visualization, particularly due to the complexity of ST data and the diversity of analysis platforms. The *SCNT* (Single-Cell, Single-Nucleus, and Spatial Transcriptomics Analysis and Visualization Tools) package was developed to address these challenges by providing an efficient and user-friendly tool for processing, analyzing, and visualizing SC and ST data.

**Results:** *SCNT* is an R-based package that integrates widely used tools such as *Seurat* and *ggplot2*, enabling seamless conversion between *Seurat* and *H5ad* formats. The package supports high-resolution spatial visualization, including customizable gene expression and clustering plots. *SCNT* also simplifies key data analysis steps, such as quality control, dimensionality reduction, and doublet detection, significantly enhancing workflow efficiency. We tested *SCNT* on publicly available PBMC dataset, Visum and Visium HD human kidney tissue data, demonstrating its effectiveness.

**Conclusions:** *SCNT* offers a valuable tool for researchers exploring SC and ST data. Its simplicity, flexibility, and powerful visualization capabilities provide a streamlined workflow for both novice and advanced users. Future developments will focus on expanding support for additional ST platforms and enhancing multi-omics data integration.

## 1. Background

With the rise of single-cell (SC) technologies, particularly spatial transcriptomics (ST), researchers are now able to explore cellular dynamics in both time and space at SC resolution [1]. Several ST technologies have gained popularity in recent years, such as 10X Genomics’ Visium and its latest version, Visium HD [2], as well as BGI’s Stereo-seq [3]. These technologies offer a rich array of options for researchers. Currently, the majority of publicly available ST datasets are based on 10X Visium, which captures transcriptomic data across a 11×11 mm^2^ area with 55 µm diameter spots. In contrast, the newly launched Visium HD offers a more detailed coverage of tissue sections by utilizing three different bin sizes: 2 µm, 8 µm, and 16 µm, thereby providing comprehensive transcriptomic data across the entire section [4].

At present, *Seurat* and *Scanpy* are the leading software tools for processing SC and ST on R and Python platforms, respectively [5, 6]. Both platforms offer a rich set of packages for the analysis of SC and ST data, making it common for users to alternate between the two. Consequently, the efficient conversion between Seurat objects and H5ad files can significantly enhance workflow efficiency. However, despite the capabilities of *Seurat* and *Scanpy*, visualizing data in a personalized manner, especially for ST data involving background tissue sections and region-specific cropping, remains challenging.

To address these limitations, we have developed a new software package designed to fill these gaps and provide a user-friendly toolbox for the analysis and visualization of SC and ST data. This tool simplifies the quality control and doublet removal processes for SC data. It also provides a straightforward method for converting *Seurat* and H5ad objects back and forth. Additionally, it allows users to easily customize the visualization of both SC and ST data according to their preferences, ensuring optimal results.

## 2. Implementation

We developed the R package *SCNT*, which facilitates the basic processing, analysis, and personalized visualization of SC and ST data. *SCNT* is fully built on the R platform (version 4.1.0 or higher) and is primarily based on Seurat (version 5.0.0 or higher) and ggplot2. The core feature of *SCNT* is its visualization capabilities, all of which are based on *ggplot2* [7]. This allows users to easily customize and modify plot styles through *ggplot2* theme parameters, enabling personalized adjustments for both SC and ST visualizations (**Figure 1**). Detailed usage instructions and function documentation for *SCNT* can be found at https://github.com/746443qjb/SCNT, which includes comprehensive examples.

**Figure 1.**
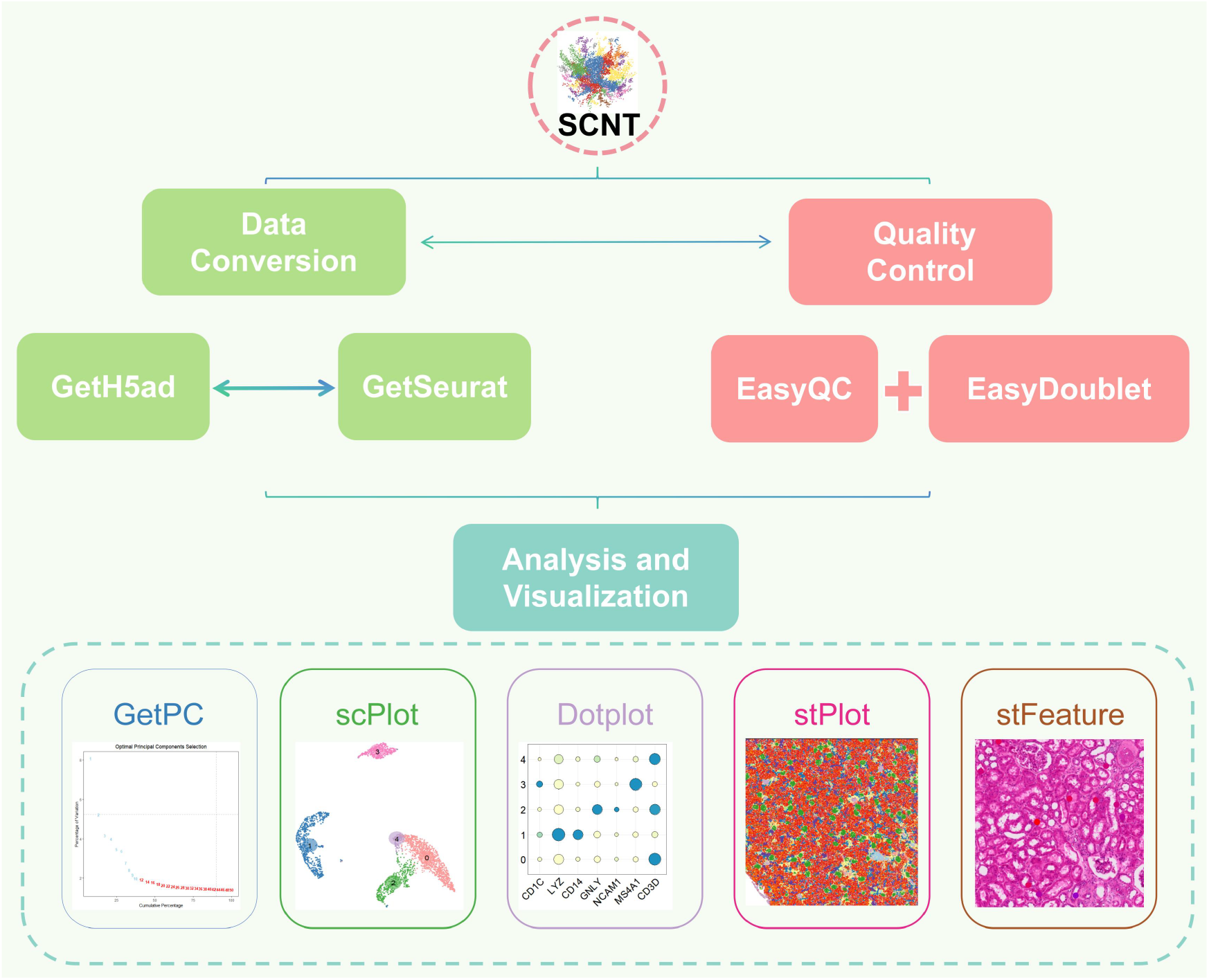
Main Functions and Workflow of SCNT. This figure illustrates the main functions and workflow of the SCNT package, including steps such as quality control, dimensionality reduction, clustering analysis, and spatial visualization. SCNT integrates multiple functional modules, providing a complete workflow from data preprocessing to final result visualization.

### 2.1 Single-Cell Data Quality Control

SCNT simplifies the standard Seurat workflow for SC data quality control, focusing primarily on the control of mitochondrial genes, erythrocyte genes, ribosomal genes, and UMI counts. Additionally, the package allows for consistent processing of both human and mouse data by setting the species parameter.

Furthermore, SCNT integrates DoubletFinder to streamline the doublet filtering process [8]. We also address the common issue of selecting the optimal number of principal components (PCs) by employing a combined method of cumulative variance contribution and the point of inflection in variance decay. Specifically, we choose the number of PCs that satisfies two criteria: (1) the cumulative variance exceeds a threshold C (e.g., 90%) and (2) the variance of the current PC is below a threshold V (e.g., 5%), with the inflection point in the variance drop exceeding 0.1. The optimal number of PCs is then determined by selecting the minimum of these two values. Where pct_*i*_ represents the variance contribution of the *i—*th PC and the cumulative variance is defined as: 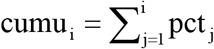. Then the best PC can be expressed as:

PC* = min (min{*i*:cumu_*i*_>*C* and pct_*i*_<*V*}),min{i>1:pct_*i*−1_-pct_*i*_>0.1}+1)

The threshold *C* and the variance of the current PC is lower than threshold *V*.

### 2.2 Conversion Between Seurat and H5ad Objects

*SCNT* uses *reticulate* to configure the Python environment within R, enabling seamless conversion between *Seurat* and *anndata* data formats [9]. This allows for the efficient transfer of raw gene expression matrices, metadata, dimensionality reduction results, and spatial coordinates for both SC and ST data. In terms of ease of use, *SCNT* offers a more user-friendly solution compared to other R packages such as *SeuratDisk* and *sceasy* [10, 11].

### 2.3 Visualization of Single-Cell and Spatial Transcriptomics Data

All visualization functions in the *SCNT* package are built upon *ggplot2*. Using functions including *dplyr, rlang*, and *GetAssayData* [12], we extract and organize gene expression and grouping information from SC and ST data, which are then visualized using *ggplot2*. The available visualization types include dimensionality reduction plots, bubble plots, scatter plots, and spatial plots.

It is important to note that, although our Spatial plot is based on *ggplot2*, we utilize the *annotation_raster* function to add background tissue images. Moreover, by applying row and column slicing on the images, we enable the visualization of cells and tissue within any specified coordinate range. This approach provides flexibility in visualizing sub-regions, unlike methods such as those used in *Scanpy*, which rely on grid-based segmentation for sub-region visualization.

## 3. Results and discussion

### 3.1 Processing and visualization of single cell data

We used the *SCNT* package to analyze and explore the peripheral blood mononuclear cell (PBMC) dataset (3000 cells) provided by 10X Genomics, as well as human kidney tissue data from Visium and Visium HD. For the PBMC dataset, we performed basic quality control using SCNT, applying the following filtering thresholds: 1) Mitochondrial (MT) genes ≤ 5%; 2) Ribosomal protein (RP) genes ≤ 5%; 3) Hemoglobin (HB) genes ≤ 1%l; 4) Feature counts ≥ 200 and ≤ 5000. After applying these filters, 2639 cells passed the quality control (**Figure 2A**). We then performed PC analysis (PCA), retaining the first 50 dimensions. Using SCNT, we found that the cumulative variance contribution and the variance decay plateaued after the 10th PC, indicating that the first 10 PCs best captured the underlying structure of the data (**Figure 2B**). Thus, we selected PC=10 as the optimal number for subsequent analysis.

**Figure 2.**
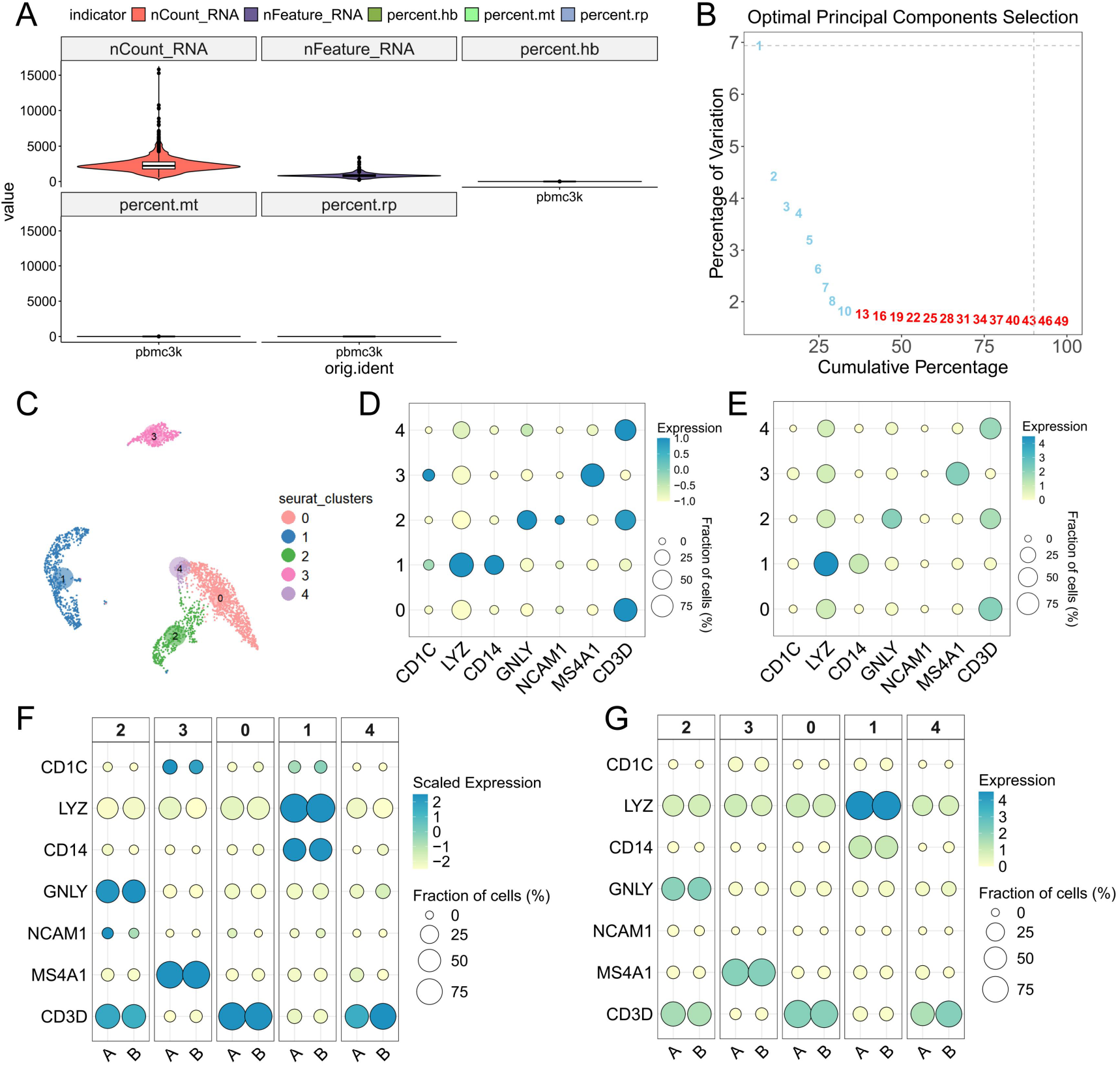
SCNT Testing on PBMC Dataset. (A) Violin plot showing the number of count RNA and feature RNA, as well as the proportions of hb (hemoglobin), red blood cells, mt (mitochondrial), rp (ribosomal) genes after quality control of the PBMC dataset using the *EasyQC* function.(B) Calculation of the optimal PCs for dimensionality reduction using the GetPC function.(C) UMAP plot of the PBMC dataset generated by *scPlot*. (D) and (E) Bubble plots displaying the expression of various markers in five cell clusters, both normalized and unnormalized. (F) and (G) Bubble plots showing the expression of the same markers in five cell clusters and across different groups (A, B), for both normalized and unnormalized data.

**Figure 3.**
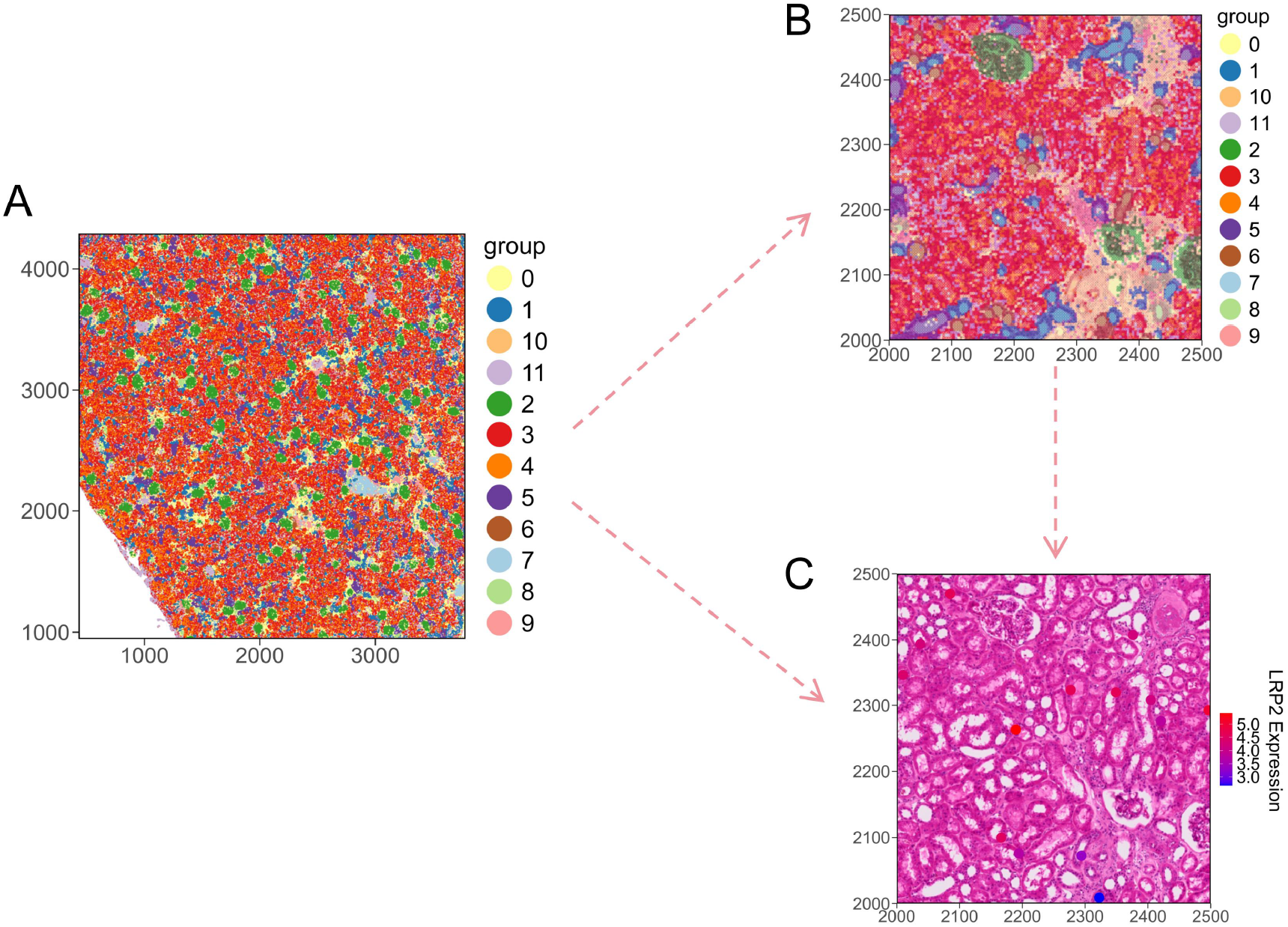
SCNT Testing on Human Kidney 10X Visium HD Dataset. (A) Spatial clustering plot based on *stPlot* showing the distribution of 12 cell clusters across the entire tissue space. (B) Spatial distribution of the 12 cell clusters within a sub-region. (C) *scFeature* plot displaying the spatial expression feature of *LRP2* within the sub-region.

*SCNT* efficiently identified 56 potential doublets and filtered them out. This simplified doublet detection process offers significant advantages over traditional methods like *DoubletFinder* and *DoubletDecon* [13], reducing the learning curve and trial-and-error for new users or those less familiar with R. Subsequently, using the 10 retained PCs, we performed clustering and UMAP dimensionality reduction on the PBMC dataset. With a resolution of 0.2, the 2583 remaining cells were divided into 5 distinct clusters (**Figure 2C**).

To demonstrate the visualization capabilities of SCNT, we selected a set of commonly used PBMC markers (*CD3D, MS4A1, NCAM1, GNLY, CD14, LYZ, CD1C*) and tested the *scDot* function [14]. The *scDot* function, similar to *Seurat’s DotPlot*, is based on *ggplot2* but retains much of the familiar functionality of *Seurat*, making it easier for users to transition between the two tools. An important feature of *SCNT* is its flexibility in gene expression visualization. While most tools display standardized gene expression values, *SCNT* allows users to visualize either normalized or raw gene expression levels in bubble plots, providing more flexibility (**Figure 2D and 2E**). As SC research grows increasingly complex, visualizing gene expression across multiple group conditions has become more important. To address this, we developed the *scMultipleDot* function. To demonstrate its capability, we randomly divided the PBMC dataset into two groups (A and B) and used it to visualize marker gene expression across different groups and cell types in a single bubble plot (**Figure 2F and 2G**).

### 3.2 Conversion Between Seurat and H5ad Objects

The *Load10X_Spatia****l*** function in the *Seurat* package, by default, only supports the reading of low-resolution tissue H&E images. However, high-resolution background images are often necessary for many analytical scenarios. To address this, we developed the *ReadST* function in the *SCNT* package, which allows users to read high-resolution H&E images and construct spatial *Seurat* objects. This function supports both *Visium* and *Visium HD* data with the same operating mode. Its parameters are similar to those in *Load10X_Spatial*, making it easy for users to quickly learn and use.

R and Python are the two most commonly used platforms for single-cell and spatial transcriptomics data processing. Many excellent analysis tools are available on both platforms, requiring researchers to switch between them. For example, *Monocle* in R is often used for pseudotime analysis [15], while *Velocyto* in Python is used for RNA velocity analysis [16]. Although packages like *SeuratDisk* and *sceasy* enable the conversion between *Seurat* and H5ad files, their workflows are often not straightforward, with many dependencies that can lead to installation and usage errors. More importantly, these tools do not support the conversion of spatial-related data, such as scaling factors and spatial coordinates, which are critical for ST analysis.

In contrast, SCNT leverages *reticulate* to integrate Python into R, allowing for simple and efficient conversion of *Seurat* objects to H5ad objects. This conversion process retains essential data from the *Seurat* object, including counts, metadata, dimensional reductions, and image parameters. Furthermore, *SCNT* also supports converting H5ad objects back into Seurat objects within R. We have tested this functionality on the processed PBMC dataset as well as human kidney tissue data from both Visium and Visium HD (**https://github.com/746443qjb/SCNT**).

### 3.3 Visualization of Spatial Transcriptomics Data

Since the advent of ST, Python has been the primary platform for visualizing ST data, with many excellent visualization packages emerging, such as *Scanpy* and *Squidpy* [17]. These tools offer a variety of functionalities to explore spatial patterns of gene expression in tissue sections. Although *Seurat* also provides basic spatial visualization capabilities, it can be challenging to achieve highly customized visualizations with its default options.

In contrast, *SCNT* leverages *ggplot*2 within R for ST data visualization. By overlaying the ST data matrix with background images, we enable easy and effective spatial clustering and gene expression visualization. Additionally, with the use of a simple image row-column slicing function, sub-region visualization becomes much more accessible. Users can input any row and column coordinate range to observe the distribution of cells and gene expression in specific sub-regions. While packages like *Squidpy* and *CellScopes* also support sub-region visualization [18], they typically do so by dividing the coordinates into fixed grid-based segments, which can be limiting for more flexible analysis.

To demonstrate *SCNT*’s functionality, we processed Visium HD data of human kidney tissue using the 10X official tutorial, including dimensionality reduction and clustering. Using *SCNT*, we visualized 12 cell clusters across the entire tissue and in specific sub-regions. By adjusting the *ggplot2* theme parameters, we were able to generate spatial cluster plots in different styles. Furthermore, we visualized the expression of genes such as *LRP*2 in particular regions [19], highlighting *SCNT*’s ability to offer detailed insights into gene localization. *SCNT* provides researchers with more flexible and customizable visualization options for spatial transcriptomics data, allowing for the creation of a variety of visual styles using *ggplot2*.

## 4. Conclusion

In this study, We introduced *SCNT*, an R package designed to simplify the processing, analysis, and visualization of SC and ST data. SCNT efficiently converts between *Seurat* and H5ad formats, enables high-resolution spatial visualization, and offers customizable gene expression plots. By streamlining key steps such as quality control, clustering, and dimensionality reduction, *SCNT* enhances the accessibility and flexibility of ST analysis. With its user-friendly features and powerful visualization capabilities, *SCNT* provides researchers with a valuable tool for exploring the spatial organization of gene expression in tissues. Future improvements may include support for additional spatial platforms and integration with multi-omics data, further advancing spatial biology research.

## Availability and requirements

Project name: SCNT (Single-Cell, Single-Nucleus, and Spatial Transcriptomics Analysis and Visualization Tools)

Project home page: https://github.com/746443qjb/SCNT

Operating system(s): Windows, MacOS and Linux.

Programming language: R

Other requirements:R ≥ 4.1.0 and Seurat ≥ 5.0.0 License: MIT License

Any restriction to use by non-academics: none

## List of abbreviations

HB: Hemoglobin
MT: Mitochondrial genes
PCA: Principal component analysis
PC: Principal component
PBMC: Peripheral blood mononuclear cell
RP: Ribosomal protein
SC: Single cell
ST: Spatial transcriptomics

## Declarations

## Ethics approval and consent to participate

Not Applicable

## Consent for publication

Not Applicable

## Availability of data and materials

The SCNT package, along with example tutorials, is available for download on GitHub at https://github.com/746443qjb/SCNT. SC and ST data from 10X Genomics can be accessed through the official 10X Genomics website or their designated data portals (https://www.10xgenomics.com/datasets).

## Authors’ contributions

Conceptualization: Jianbo Qing. Methodology: Jianbo Qing. Investigation: Jianbo Qing. Validation: Jianbo Qing. and Yafeng Li. Visualization: Jianbo Qing. Formal analysis: Jianbo Qing. Funding acquisition: Junnan Wu. Project administration: Junnan Wu. Supervision: Junnan Wu. Writing–original draft: Jianbo Qing. Writing–review & editing: Jialu Wu.

## Acknowledgements

We would like to thank those who provided guidance and support throughout the course of this work.

